# Toward Holistic Solutions to Undernutrition: A Comprehensive Preclinical Protocol for Novel Ready-to-Use Therapeutic Foods Testing

**DOI:** 10.1101/2024.11.28.625813

**Authors:** Laura Nguoron Utume, Jerome Nicolas Janssen, Peter Akomo, Daisy Shaw, William Joseph Samuel Edwards, Molly Muleya, Charoula Konstantia Nikolaou, Aviv Schneider, Andrew Westby, Ram Reifen, Anastasios D. Tsaousis, Efrat Monsonego-Ornan, Aurélie Bechoff

**Affiliations:** Natural Resources Institute, University of Greenwich, Central Avenue, Chatham, ME4 4TB, UK; Department of Biological Sciences, Benue State University, Makurdi, PMB 102119, Nigeria; School of Natural Sciences, University of Kent, Canterbury, CT2 7NH, UK; Department of Biochemistry and Nutrition, The Hebrew University of Jerusalem, Robert H. Smith Faculty of Agriculture, Food and Environment, Rehovot, Israel; Valid Nutrition, Cuibín Farm, Derry Duff, Bantry, Co., Cork, Republic of Ireland; University of Nairobi, Department of Food Science, Nutrition and Technology, University of Nairobi; School of Biosciences, Division of Food, Nutrition and Dietetics, University of Nottingham, Sutton Bonington Campus, Loughborough, Leicestershire, LE12 5RD, UK; University of Greenwich, Park Row, Greenwich, SE10 9LS, UK

**Author notes:** Shared first authors (Laura Nguoron Utume, Jerome Nicolas Janssen). Corresponding authors: Dr. Anastasios D. Tsaousis Dr. Aurelie Bechoff Prof. Efrat Monsonego-Ornan.

**Keywords:** RUTF, Malnutrition, Bone, Metabolomics, Microbiome

## Abstract

Severe acute malnutrition (SAM) affects over 45 million children under five worldwide. Ready-to-Use Therapeutic Foods (RUTFs), the cornerstone of treatment, reach only 30% of children in need due to high production costs and reliance on imports. RUTF formulations developed from local ingredients have shown varying efficacy levels focusing primarily on weight gain, overlooking critical factors like micronutrient status, skeletal development, and gut health. The presented study establishes a comprehensive protocol for preclinical testing of novel RUTF formulations using a validated juvenile rat model replicating key features of malnutrition in children. Beyond the standard anthropometric measurements (body weight, length, and food consumption), bone quality is assessed through micro-computed tomography and mechanical testing, providing insights into microstructure and functionality beyond standard anthropometric measurements. Concurrently, gut microbiome dynamics are analysed using 16S rRNA sequencing coupled with metabolomic profiling of stool and serum to monitor gut health and systemic recovery.By addressing critical gaps in traditional RUTF evaluations and emphasising locally sourced, cost-effective formulations, this protocol offers a transformative framework for developing accessible and impactful malnutrition treatments. The findings generated through this multidisciplinary approach will guide RUTFs optimisation, strengthen their scientific foundation, and pave the way for evidence-based clinical trials to improve global malnutrition treatment strategies.

## Introduction

Globally, 45 million children under the age of five years suffer from acute malnutrition (AM), with one-third of them affected by severe acute malnutrition (SAM). Children with SAM are 9–12 times more likely to die than their healthy peers^1^. Ready-to-Use Therapeutic Food (RUTF) is widely used to combat malnutrition; however, its high production costs and expensive imports —primarily due to manufacturing in developed countries—limit its coverage to only 30 % of children with SAM^2^. Codex guidelines^3^ now permit the use of alternative RUTF formulations made from local foods to address the shortage. These alternatives can lower costs, support local economies^4,5^, and improve recovery outcomes traditionally assessed by weight gain. However, critical factors such as micronutrient status^6^, linear growth, and gut health^7^, which are essential for long-term intervention success in reducing the risk of metabolic diseases later in life, are often overlooked^8^.

Recent RUTF designs have incorporated knowledge of gut microbiome maturation to address the need for not only cost-effective RUTFs but also a more holistic approach to evaluating their effectiveness. These approaches have shown promising results, including improvements in anthropometric measurements and plasma protein mediators of bone growth, gut repair, and inflammation^9–11^. In this study, we establish a protocol to conduct preclinical trials on rats to develop and assess alternative RUTFs for treating malnutrition.

### RUTF design and animal models

Preclinical research heavily relies on animal experimental models due to their ability to replicate complex biological systems and disease processes.^12^. For example, they can simulate dietary interventions, disease processes, and therapeutic testing within a complex and integrated biological system^12,13^. However, a significant challenge in animal modelling is its translatability, as successful outcomes in animal experiments do not always reliably predict results in human subjects^12–14^.

Rodents are now the species of choice for biomedical research due to their ease of handling, breeding, and maintenance and their anatomical, physiological, and genetic similarities to humans^15,16^. Additionally, rats are particularly suitable for dietary research since their nutritional requirements and food choices resemble those of humans. Their protein digestibility values also closely align with those of humans^17^, and they share a comparable amino acid requirement^18^. This makes them an excellent model for evaluating the development of novel RUTFs.

In formulating a novel RUTF, protein quality is initially assessed using scoring metrics such as the Digestible Indispensable Amino Acid Score (DIAAS) or the traditional Protein Digestibility-Corrected Amino Acid Score (PDCAAS). However, these metrics do not always correspond to the growth patterns observed in rats^19^. For example, plant-based protein sources often contain anti-nutritional factors like trypsin inhibitors, phytates and tannins^20^, which impair protein digestion and elevate protein requirements^21^, necessitating careful consideration of these factors during RUTF development. Bone quality is also affected by micronutrients such as calcium, phosphorus, and vitamin D, zinc, copper, iron^22–26^ and, therefore, must be sufficiently available and provided during skeletal growth.

### Monitoring skeletal growth

In the context of RUTF design, current assessments of malnutrition and its interventions often rely primarily on overall weight measurements. However, a recent study on rats fed a 20% control protein diet and a 10% deficient protein diet demonstrated that, despite no significant differences in final body weight between the two groups, the mechanical behaviour of femoral bones differed significantly in the 10% group^19^. This highlights the need for more detailed assessments of bone quality during nutritional studies^27,28^.

To address this, we employ various techniques for in-depth bone analysis while continuous weight, length, and food consumption measurements conducted throughout the experiment monitor growth trajectories. Following a 6-week experimental period, bone samples will be collected for microCT analysis and three-point bending assays, providing insights into bone microarchitecture and mechanical properties.

### Multi-Omics approach to evaluate the host’s health

Gut health and microbial composition play a critical role in treating malnutrition. Consequently, RUTF formulations should aim to support microbiome maturation and restore normal gut function. Microbiota-directed complementary foods have shown promise in increasing the abundance of key bacterial species involved in gut community development ^7,10,11^.

To evaluate these effects, rigorous microbiome analyses will be conducted on stool samples collected multiple times during the experiment and from the cecum after sacrifice. These analyses will include diversity metrics and differential bacterial abundance studies. Simultaneously, metabolomic profiling will be performed on the same samples to assess gut function, intervention success, and microbial activity. Previous research has established links between the microbiome and metabolome, identifying associations between specific metabolites and gut enterotypes^29^. Additionally, caecal samples will provide a more localised view of microbial populations and metabolic activities within the intestinal tract, complementing stool-based analyses.

This pilot study aims to establish a comprehensive experimental framework for evaluating malnutrition and testing novel interventions in a rat model, overcoming the limitations of traditional methods. By incorporating detailed bone analysis, microbiome profiling, and metabolomic studies, this approach provides a multidimensional perspective on recovery, enabling more accurate predictions of long-term outcomes. Building on the findings from this pilot study, this protocol will be expanded to include additional RUTF groups, with dietary interventions introduced 3 weeks into the experimental timeline to assess their impact on growth and health restoration.

## Material and Methods

### Preclinical experiment using animal (rat) models

This preclinical pilot study utilised female Sprague Dawley (SD) rats to model underweight conditions, including wasting and stunting, observed in children aged 6–59 months. The experiment aimed to mimic malnutrition and assessed treatment effects, including potential recovery. Conducted over 6 weeks, the study was carried out under controlled environmental conditions (23 ± 1°C) with a 12-hour light/dark cycle (8 a.m. – 8 p.m.) in a Non-SPF (Conventional) animal facility at the Robert H. Smith Faculty of Agriculture, Food and Environmental Sciences, Hebrew University of Jerusalem, Israel (HUJI). Ethical approval was obtained from the HUJI Authority for Biological and Biomedical Models (ABBMS) (Ethics Number AG-22-16962-3) under the established guidelines for the care and use of laboratory animals.

Three-week-old female SD rats, weaned and acquired from Harlan Laboratories (Rehovot, Israel), were weighed upon arrival and randomly assigned to treatment groups of eight animals each, housed in two cages (four rats per cage) to create two technical replicates per group. Individual rats were identified by ear piercing. The study included a continuous standard diet group (CT) and a protein- and micronutrient-deficient diet group (MN), both serving as baseline groups for a potential comparison with novel RUTF formulations not included here. Before the experiment, rats underwent a three-day acclimation period. Throughout the study, all rats had ad libitum access to food (including standard chow, balanced and protein-deficient and treatment diets) and water.

During the first 3 weeks, an accelerated growth phase was simulated. The CT (control and potential RUTF groups) received the AIN-93G-recommended diet for growing rodents. In contrast, the MN (control and potential RUTF groups) were fed a protein- and micronutrient-deficient diet to mimic malnutrition. For the following 3 weeks, the CT control and MN control groups continued their respective diets, while potential CT and MN groups would transition to either RUTF diets as a positive control (CT-RUTF) or the RUTF intervention itself (MN-RUTF) (**Figure 1**).

**Figure 1:**
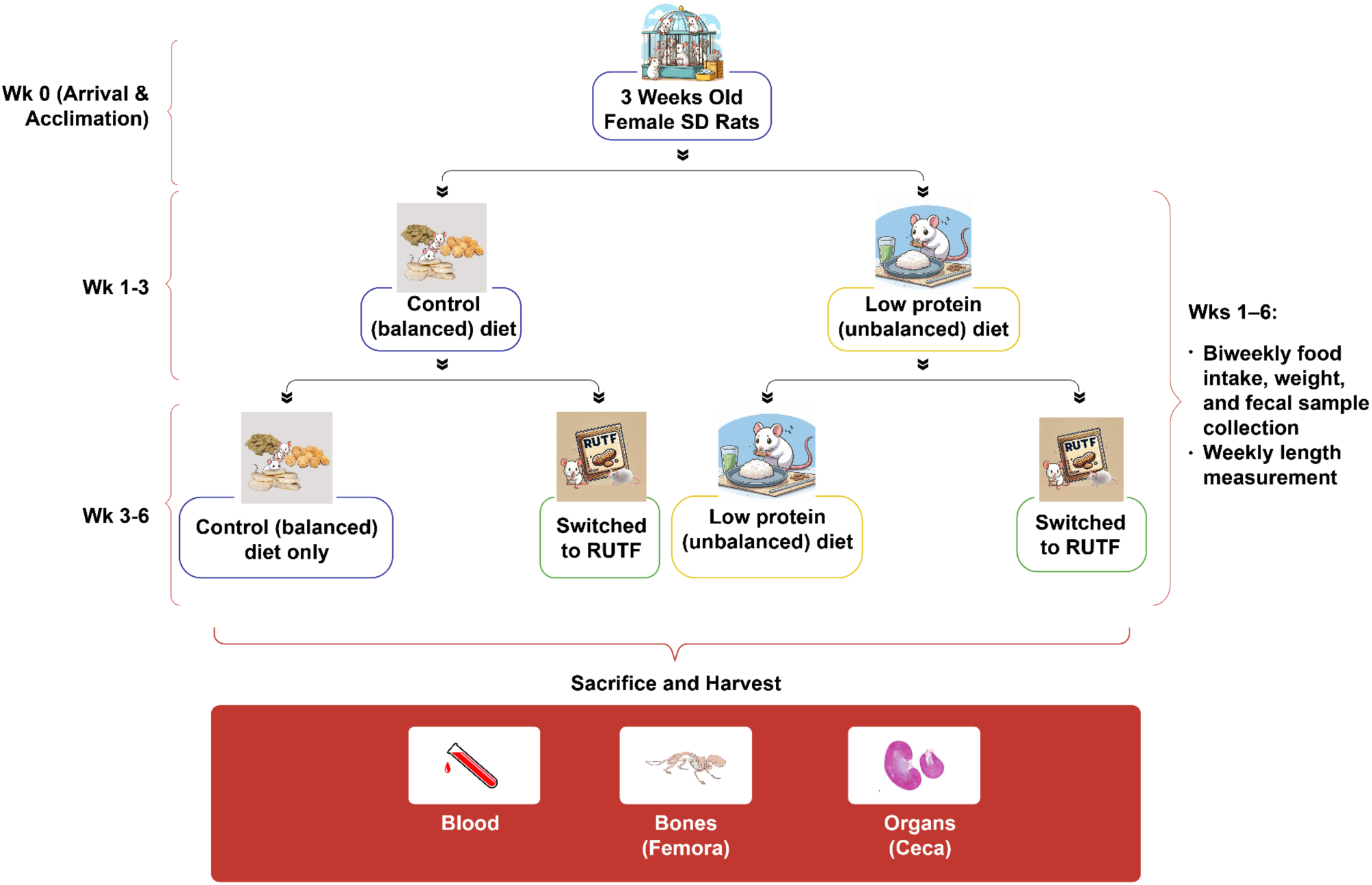
Schematic Diagram of Preclinical Experimental Set-up and Sample Collection

Throughout the experimental period, food consumption (g) and body weight (g) were measured twice weekly, while body length (cm), measured from the nose to the tail’s tip, was recorded weekly. Faecal samples were collected twice weekly in duplicate, with one sample stored in RNA/DNA Shield for 16S-Sequencing and the other in 100% methanol for metabolomics. Rats were monitored regularly for physical and behavioural changes.

At the end of the 6 weeks, all rats were anaesthetised with Isoflurane USP 100% for final measurements and sample collection. Weight (g) and length (cm) were recorded under anaesthesia. Blood samples were collected via cardiac withdrawal and centrifuged at 2000 x g for 20 minutes at 4 °C. The serum was transferred into sterile two ml tubes and stored at -80 °C for metabolome analysis. Cecal samples were collected in sterile five ml tubes, flash-frozen in liquid nitrogen for 5 to 10 minutes, and stored at -80°C for microbiome sequencing. Femurs were harvested, cleaned of soft tissue, wrapped in gauze soaked in phosphate-buffered saline (PBSx1), and preserved at -20°C for mechanical and micro-CT analysis. All sample handling was performed under aseptic conditions.

### Control diet composition and preparation

The control balanced diet (CT) was formulated at HUJI following the AIN-93G guidelines for growing rodents. The diet consisted of 16 % fat, 63.5 % carbohydrates, and 20.5 % protein, supplemented with a full complement of multi-minerals and multivitamins (**Table 1**). The protein-deficient diet (MN) was also based on AIN-93G recommendations, with modification, containing only 5% protein, with 50% of the recommended multimineral and multivitamin content and 25 % fat, 70 % carbohydrates (**Table 1**). All ingredients, sourced commercially per AIN-93G specifications, were weighed, homogenised, and mixed with water to form a firm dough. The dough was moulded into medium-sized patties and air-dried in an extractor hood for 42 hours at ambient temperature. Once dried, the patties were weighed and stored at -20 °C until use.

**Table 1:** Ingredient composition of CT and MN diets.

| Product/Ingredient | Control diet<br>(CT)(g/kg diet) | Protein and<br>micronutrients<br>deficient diet<br>(MN)(g/kg diet) |
| --- | --- | --- |
| Cornstarch | 397 | 438.7 |
| Dextrinized Cornstarch | 132 | 145.7 |
| Sucrose | 100 | 110 |
| Fibre | 50 | 50 |
| Soybean Oil | 70 | 120 |
| Casein ($\geq 85\%$ Protein) | 200 | 50 |
| L-Cystine | 3 | 3 |
| Mineral mix (AIN-93G-MX) | 35 | 17.5 |
| Vitamin mix (AIN-93G-VX) | 10 | 5 |
| Choline bitartrate (41.1% Choline) | 2.5 | 1.34 |
| Tert-Butylhydroquinone | 0.014 | 0.015 |

### RUTF design and analysis

RUTF formulations were developed theoretically using linear programming^30^ to align with the 2022 UNICEF RUTF Specifications and Codex requirements^3^. These formulations were tested for nutritional conformance to ensure they met established standards. Prototype batches (2.5 kg per formulation) were produced under sanitary conditions and evaluated for nutritional compliance. Subsequently, larger batches (20 kg) were produced and checked for aflatoxin levels, microbiological safety, and amino acid bioavailability using the Digestible Indispensable Amino Acid Score (DIAAS). For comparison, a commercially available standard RUTF, such as one from Nuflower Foods and Nutrition Private Limited (Haryana, India), can be procured. Its nutritional composition will be verified and used as a control to benchmark the novel RUTF formulations.

### Bone structure analysis by computed microtomography (Micro-CT)

Femora were scanned using a Skyscan 1272 X-ray micro-CT system (Bruker) at 50 kV X-ray tube voltages and 800 μA with a 0.25 mm aluminum filter, 4000 ms exposure time, and a resolution of 15 μm. Each sample yielded 900 projection images through a 180° rotation with 0.4° steps, averaged over two frames. Image reconstruction was conducted with NRecon software (Skyscan), and morphometric analysis was performed using CTAn software (Skyscan) per established guidelines for bone microstructure analysis^31,32^. For cortical bone analysis, a region comprising 150 slices was selected with a global grayscale threshold of 66-255. For the trabecular region, 150 slices were analyzed with adaptive grayscale thresholds of 49-255. Reconstructed images were rendered in 2D and 3D using Amira software (v.6.4, FEI, Hillsboro, OR, USA). Cortical bone parameters included total cross-sectional area (Tt.Ar), cortical area (Ct.Ar), cortical area fraction (Ct.Ar/Tt.Ar), cortical thickness (Ct.Th), medullary area (Ma.Ar), and bone mineral density (BMD). Trabecular bone parameters included bone volume fraction (BV/TV), trabecular number (Tb.N), trabecular thickness (Tb.Th), and trabecular separation (Tb.Sp). Femur length was measured using SkyScan software^13^. Wilcoxon rank-sum test was used to assess significance.

### Bone mechanical analysis by three-point bending

The mechanical strength of the femur from each rat was evaluated by three-point bending using an Instron 3345 material testing machine equipped with a custom saline chamber^33^. A support span of 12 mm was used to position the mid-diaphyseal region of each femur. A preload of 0.1 N was applied to stabilise the bone, and force-displacement data were captured at 10 Hz via Instron software (BlueHill). From the force-displacement curves, bone stiffness, yield point, fracture load, maximum load, and total energy to fracture (AUC) were calculated. Bone stiffness was determined from the linear region of the curve, incorporating both geometric and material properties. Young’s modulus was also calculated from the stress-strain curve to assess material stiffness independently of bone length and architecture^34,35^. Wilcoxon rank-sum test was used to determine significance.

### Microbiome analysis

Defrosted cecum contents (0.25 g) were extracted using the Invitrogen PureLink™ Microbiome DNA Purification Kit according to the manufacturer’s instructions. The extracted DNA was sent to Novogene for 16S rRNA sequencing of the V3-V4 region using the 341F-806R primers. The DADA2^36^ amplicon workflow (v1.24) was used to process forward and reverse reads. Taxonomy was classified using DECIPHER (v2.24) and the RDP (v18) training set. The phyloseq package (v.1.40) was used to plot the distribution of the bacterial taxa at the phylum level, and only phyla passing the minimum frequency of 1 % were included. α- and β-diversity were assessed using the vegan (v2.6.4) package. α-diversity was calculated using the Shannon index and tested with the Wilcoxon rank sum test. After checking the homogeneity of group dispersion (p > 0.05), β-diversity was tested using PERMANOVA. Differential abundance analysis of bacteria was performed with ANCOMBC2 of the ANCOMBC^37^ package (v1.6.4) and standard parameters with a prevalence cut of 0.33 and FDR adjustment.

### Metabolomics

For metabolite extraction from serum, 200 µl of serum was added to 200 µl of MeOH and vortexed for 30 seconds. After incubation at -20 °C for 20 minutes, samples were centrifuged at 15,000 x g and 2 °C for 30 minutes. The supernatant was collected and lyophilised via freeze-drying. For metabolite extraction from stool and cecum, 0.4 - 0.6 g of cecal samples were added to 200 mg of 0.4 mm diameter glass beads and 1 ml of 80% ethanol (pre-heated to 80°C), before vortexing for 30 seconds. Samples were incubated at 80°C for three minutes, then vortexed for another 20 seconds. Glass beads were removed via centrifugation at 16,000 x g for ten minutes, at room temperature, splitting supernatant between two 2 ml tubes. This step is crucial so that the tubes are not full regarding lyophilisation. Samples were lyophilised via freeze-drying. To prepare the NMR buffer, 538 mg NaH_2_PO_4_.H_2_O, 866.2 mg Na_2_HPO_4_ and 30 ml D_2_O were combined, and the pH* was adjusted to 7.4 using NaOD (D_2_O) or DCl (D_2_O). 1 M sodium azide (NaN_3_) solution was prepared by combining 65.01 g NaN_3_ with 1 ml buffer solution in a fume hood. 150 µl of this sodium azide solution and 112.2 mg sodium trimethylsilylpropanesulfonate (DSS) were added to 30 ml of NMR buffer to form the final NMR solution. Lyophilised metabolite extraction desiccates were resuspended in 700 µl of NMR solution, and 220 µl was added to 3 mm NMR tubes.

One-dimensional (1D) 1H NMR spectroscopy was performed on a 600 MHz AVANCE III spectrometer equipped with a QCI-P cryoprobe (Bruker) at 298 K with a transmitter frequency of 600.05 MHz. Due to its high salt content, the sample had a volume of 220 µl and was measured in a 3 mm tube. The spectrometer was locked to D2O (5% v/v), with 0.1% w/v DSS as a chemical shift reference. Tuning and shimming were carried out automatically for each sample, as was the 90° pulse calibration. The receiver gain was limited to a maximum value of 128. Data used for metabolite concentration analysis were drawn from Carr-Purcell-Meiboom-Gill (CPMG) spectra with a CPMG period of 76.8 ms, consisting of 128 CPMG cycles used to suppress protein signals^38,39^. The CPMG spectra were measured using 128 scans and 16 dummy scans with a spectral width of 16.02 ppm (9615.38 Hz), giving an acquisition time of 1.70 seconds, a relaxation delay of three seconds was used, and the data size was 32,768 points giving a total recycle time of 4.7 seconds. For all experiments, the water resonance was optimised (automatically) for maximum suppression (o1p was ∼4.699 ppm). A purge pulse sequence of pulses with field strengths of 49.96 Hz and 4 1 ms smoothed square gradient pulses of amplitude -13.17, 52.68, -17.13 and 68.52% were applied for water suppression without amplitude modulation.

Bruker Topspin 3.6.3 was used to process the raw spectral files, including phasing and baseline correction. Chenomx Suite 8.4 was used to annotate processed spectra using the Profiler tool. Acquired spectra were fit to the spectra of the 600 Hz Chenomx reference database and manually adjusted to fit as closely as possible. Once fitted, metabolites and their relative concentrations were exported to CSV files. For statistical analysis of metabolomics data, the web server MetaboAnalyst 6.0 was used. Relative metabolite concentrations were uploaded to the Statistical Analysis [one factor] section. Data were filtered with a 10% interquartile range (IQR) variance filter. Data were normalized using normalisation by median and scaled by auto-scaling. Chemometric analyses were performed for both multi-group analysis and comparison of two groups. 2D scores plots of unsupervised principal component analysis (PCA) and supervised partial least squares discriminant analysis (PLS-DA) were generated. For PCA, permutational analysis of variance (PERMANOVA) was performed, with a significance threshold of p < 0.05. For PLS-DA, a 10-fold cross-validation and a permutation test set to 1000 permutations were performed. Metabolites crucial to group differentiation were established using variable importance in projection (VIP) plots and metabolites with VIP scores >1.2 were considered significant. To compare two individual treatment groups, univariate analysis was performed with log2(FC) > 1 and p value < 0.05. From this, volcano plots were produced and annotated. Comparisons between the two groups were made, and metabolites with significant p values were selected for pathway analysis. Samples were median-normalised and auto-scaled before being compared to the KEGG pathway library for *Escherichia coli* K-12 MG1655 and displayed as a scatter plot.

For biomarker analysis, statistically significant metabolites between two comparison groups were uploaded, and data were normalised by media and scaled by auto-scaling. Classical univariate receiver operating characteristic (ROC) curve analyses and area under the curve (AUC) were performed to assess the sensitivity and specificity of metabolites in predicting groups. The area under ROC curve (AUROC) values >0.8 were considered as good distinguishers of different treatment groups.

## Expected results

The proposed study addresses the limitations of previous malnutrition research by integrating bone, microbiome, and metabolomic analyses. Using exemplary analyses based on the CT (control) and MN (malnourished) groups, we provide an overview of the anticipated outcomes at week 6 of the experiment, excluding the impact of RUTF intervention. It is expected that RUTF intervention will shift the results of the MN group toward those observed in the CT group.

### Before the experiment

The basic protocol includes only the CT and MN groups for a representative baseline analysis. When testing novel RUTFs, two additional groups per RUTF must be included: CT-RUTF and MN-RUTF. After 3 weeks of the experiment, these groups will switch from their respective CT or MN diets to the RUTF. A commercially available RUTF should also be included to compare the efficacy of the novel formulations.

Novel RUTFs, designed through methods such as linear programming^30^, should undergo prototype production. These formulations must meet the 2022 UNICEF RUTF Specifications and Codex requirements^3^. Quality checks include aflatoxin levels, microbiological safety, and amino acid bioavailability, assessed using methods like the Digestible Indispensable Amino Acid Score (DIAAS).

### Throughout the experiment

During the experiment, stool samples will be collected biweekly for time-series analyses of the microbiome and metabolome. Length, weight, and diet consumption will be monitored continuously to evaluate growth patterns and dietary intake. Representative analysis of the CT and MN groups has already shown significant differences in length and weight after the experiment’s initiation (**Figure 2A and 2B**). These metrics will be used to determine the speed and effectiveness of RUTF intervention in mitigating malnutrition symptoms. The MN group consumed fewer calories (kcal) than the CT group despite the MN diet having a higher caloric density (3.5 vs. 3.2 kcal/g respectively) (**Figure 2C**). Protein efficiency analysis revealed that CT animals gained less weight per unit of protein consumed, suggesting that their surplus energy was partially directed toward energy metabolism rather than growth (**Figure 2D**).

**Figure 2:**
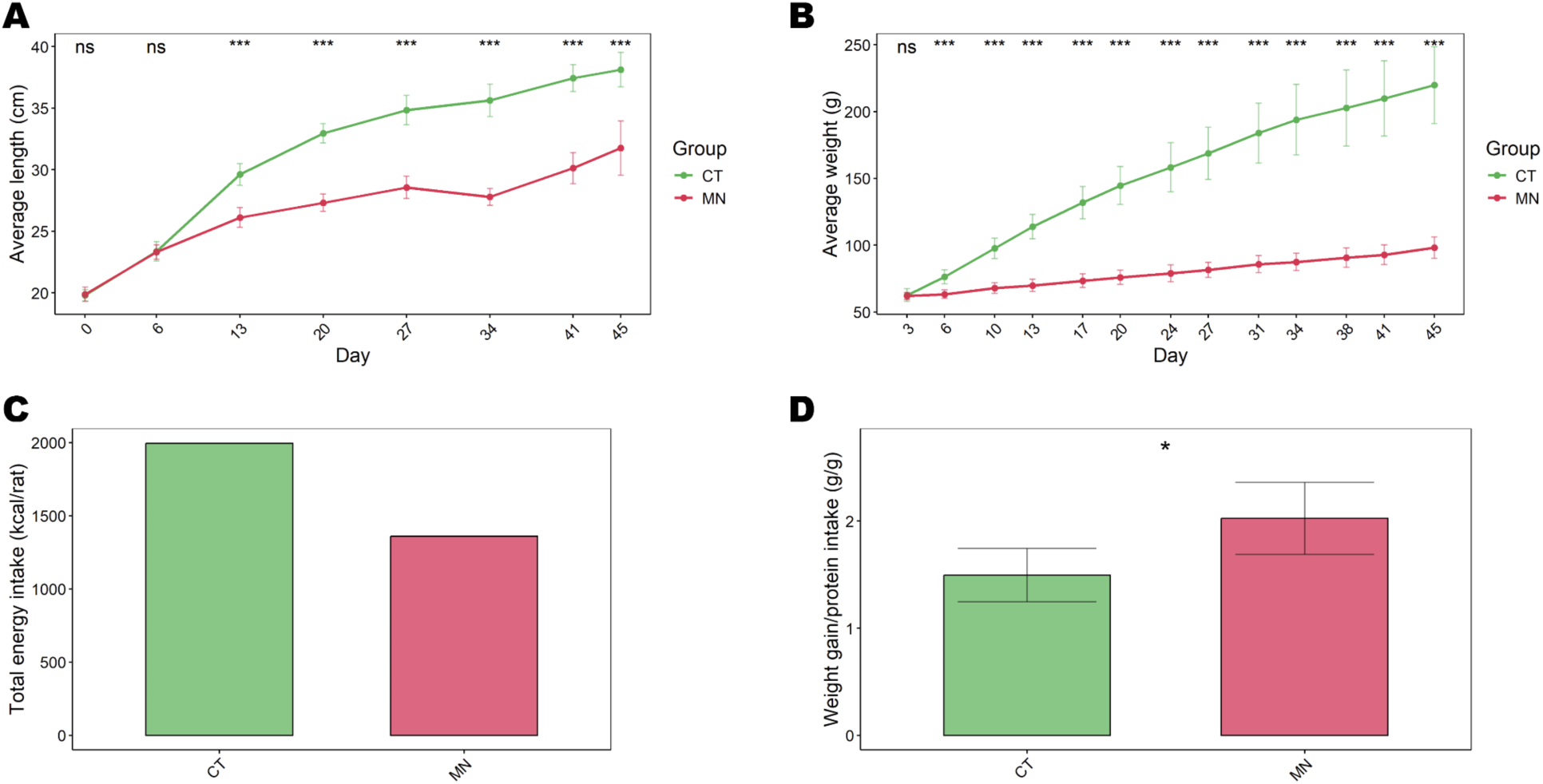
Growth parameters monitored during the experiment. **(A)** Longitudinal growth (cm/rat), **(B)** weight gain (g/rat), **(C)** total caloric intake (kcal/rat) and (**D**) protein efficiency of CT and MN. ns = not significant, * p < 0.05, *** p < 0.001

### After sacrifice

Following sacrifice at week 6, an in-depth bone analysis will be performed^40^. The femur (**Figure 3A**) will be collected for detailed assessments. Measurements will include femur length (**Figure 3B**) and cortical and trabecular bone characteristics using microCT. Parameters such as bone volume and mineralisation will be quantified to evaluate structural integrity (**Figure 3C** and **3D**). A three-point bending assay will be conducted on the femur to assess functional bone mechanics. This test is particularly significant, as bone strength and flexibility are expected to be severely compromised by malnutrition.

**Figure 3:**
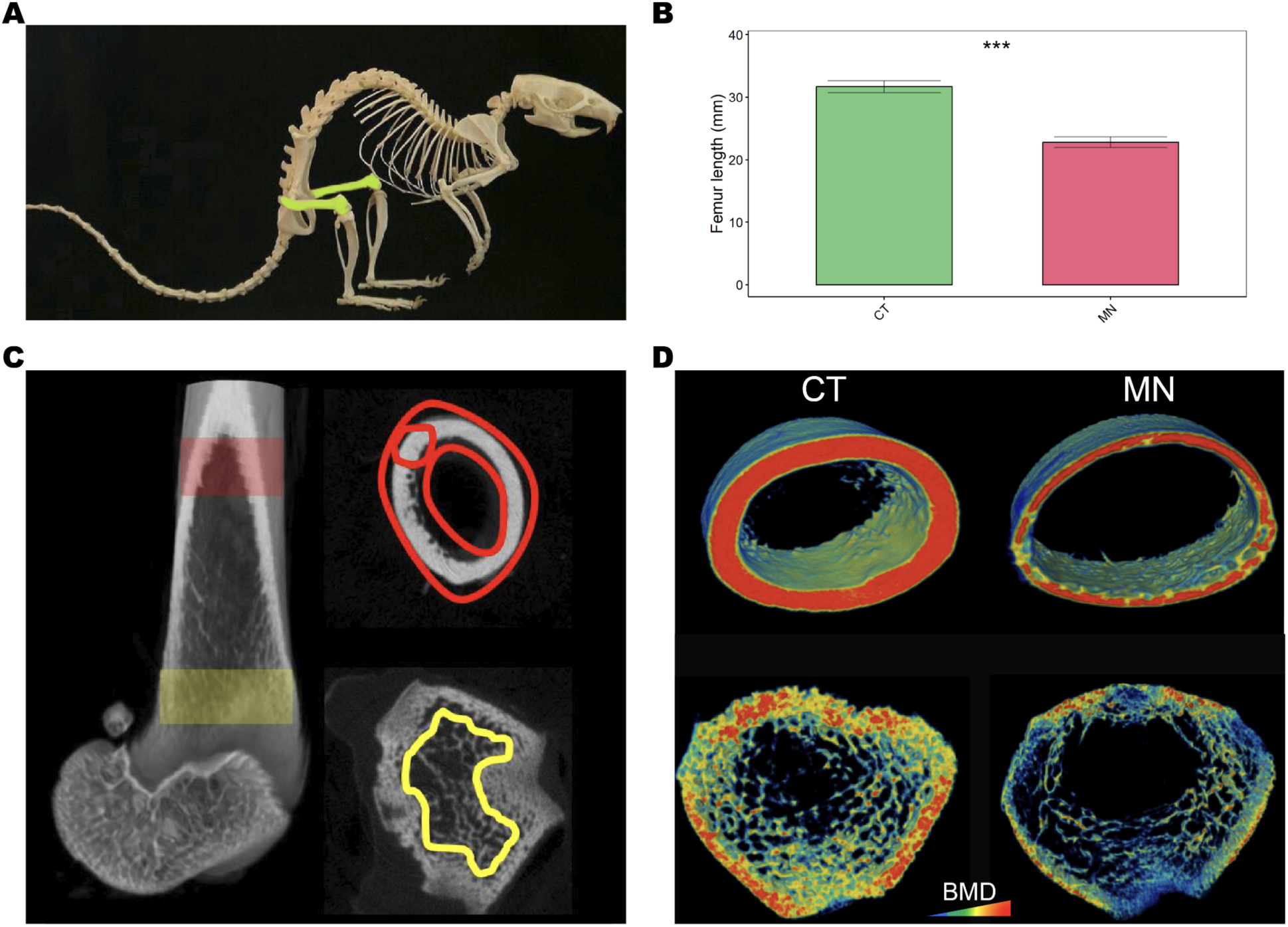
Overview of the bone analysis after sacrifice on week 6. **(A)** Skeletal scheme highlighting the femur used for the analysis and (**B**) femur length comparison of CT and MN. (**C**) Schematic overview of cortical and trabecular bone. (**D**) Comparison of the CT and MN groups’ cortical bone (top) and trabecular bone (down). Colors indicate BMD scale.

Next, a microbiome analysis of the caecum will be conducted to evaluate gut health. Preliminary findings from a representative analysis of the CT and MN groups revealed a higher relative abundance of the Actinobacteria phylum in some MN samples compared to the CT group (**Figure 4A**). However, alpha diversity, as measured by the Shannon index, did not differ significantly between the groups (**Figure 4B**). Beta diversity analysis, on the other hand, indicated a significantly distinct microbiome composition between the CT and MN groups (p = 0.0002; **Figure 4C**). Differential abundance analysis identified four genera—*Ruminococcus*, *Eubacterium*, an unclassified *Clostridiales Incertae Sedis XIII*, and *Monoglobus*—as being significantly reduced in the MN group compared to the CT group (**Figure 4D**). Furthermore, structural zero analysis using ANCOM-BC2 revealed that *Lactobacillus* was exclusively present in the CT group. In contrast, *Faecalibaculum* was uniquely detected in the MN group (data not shown).

**Figure 4:**
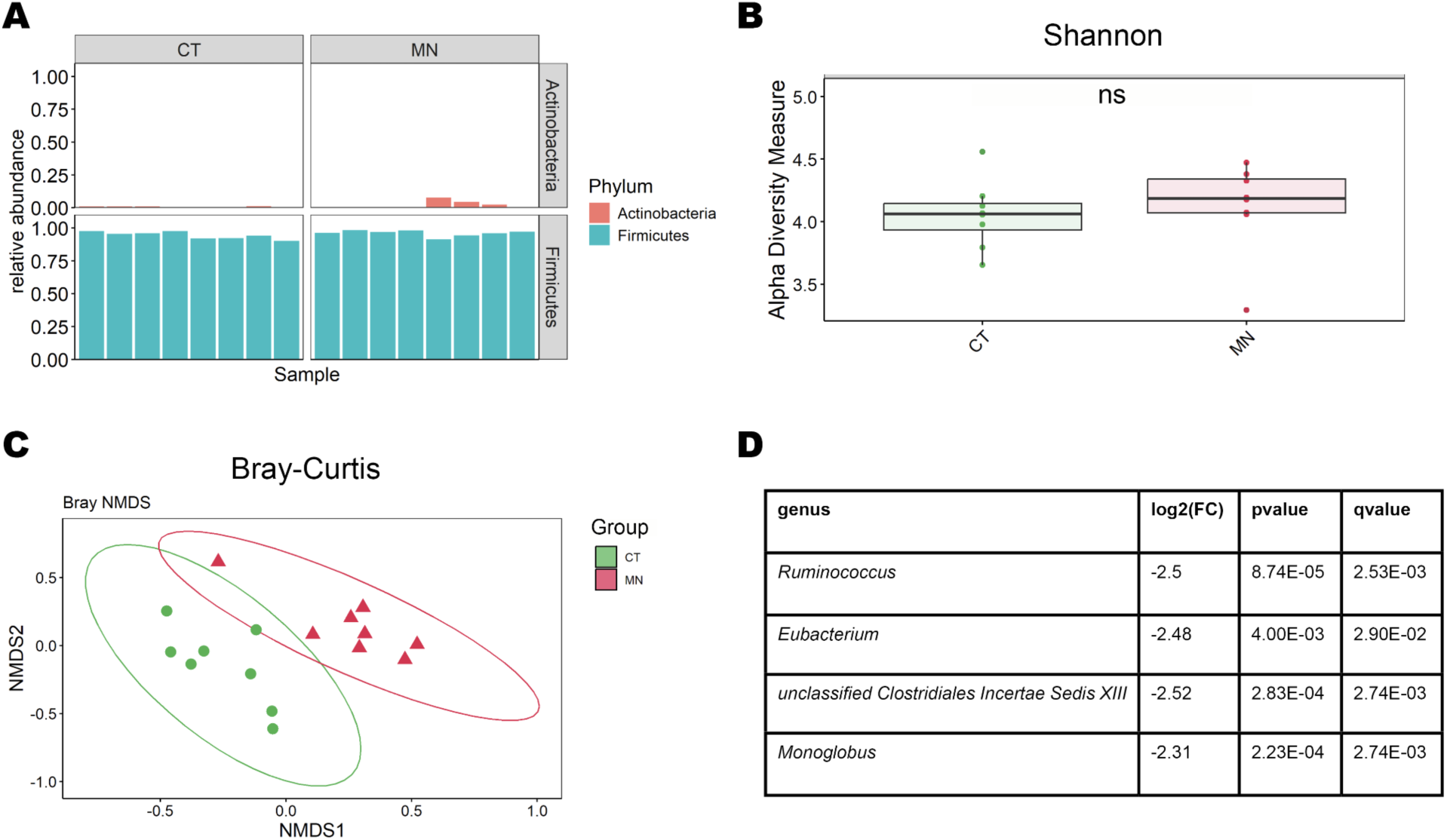
Microbiome analysis of CT and MN caecal samples. (**A**) Relative abundance of phyla surpassing a threshold of >1% per sample. (**B**) Alpha diversity as measured by Shannon index and tested by Wilcoxon rank sum test (ns = not significant). (**C**) Beta diversity as shown by NMDS plot of Bray Curtis dissimilarity. After testing for homogeneity of dispersion, PERMANOVA revealed a significant difference (p = 0.0002) of microbiome composition between CT and MN. (**D**) Differentially abundant bacteria detected by ANCOM-BC2 with p.adj < 0.05 and passing a pseudo-count addition test.

Gut functionality and microbiome health will be further investigated using metabolomics, focusing on blood serum and, as demonstrated here, the caecum of the CT and MN groups. PERMANOVA analysis of the caecal metabolome revealed a significant difference between the CT and MN groups (p = 0.016; **Figure 5A**). A total of 55 metabolites were identified with a VIP score > 1.2, making them strong discriminators between the CT and MN groups. Among these, metabolites such as erythritol and acetate were more abundant in the CT group, while others like malonate and histamine were more abundant in the MN group (**Figure 5B** and **C**). Further analyses, such as PLS-DA and ROC curve evaluations, can add to the identification of key discriminatory metabolites, while pathway analysis may provide a comprehensive understanding by integrating data across multiple metabolites.

**Figure 5:**
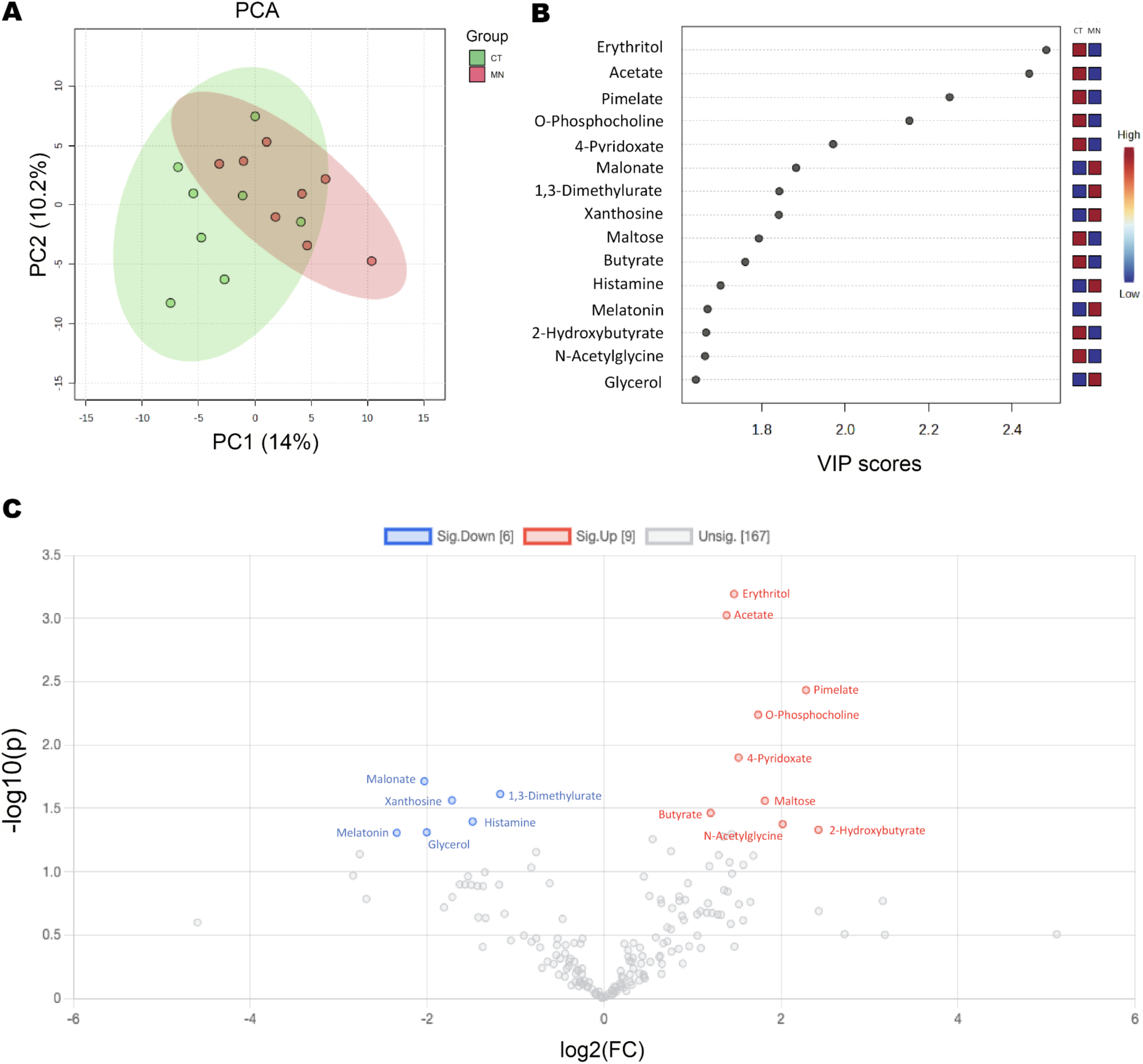
Metabolome analysis of CT and MN caecal samples using MetaboAnalyst 6.0 (www.metaboanalyst.ca). (**A**) PCA plot showed overlapping clusters of CT and MN samples (PERMANOVA: p = 0.016). (**B**) Top 15 metabolites for discriminating between healthy and malnourished groups. (**C**) Differential analysis of metabolites with a log2(FC) > 1.0 and p < 0.05. Metabolites in red are more abundant in CT, whereas metabolites in blue are more abundant in MN.

## Discussion

This pilot study outlines a comprehensive protocol for evaluating the efficacy of novel Ready-to-Use Therapeutic Foods (RUTFs) in treating severe acute malnutrition (SAM) using a preclinical rat model. By integrating advanced bone, microbiome, and metabolomic analyses, the framework addresses critical gaps in traditional approaches, which often rely solely on weight gain as a recovery metric. This multidimensional approach ensures a more holistic understanding of malnutrition and its recovery trajectory, offering robust data to inform clinical interventions.

Continuous measurements of weight, length, diet consumption and protein efficiency throughout the experiment will determine if the RUTF provides sufficient energy and protein to support optimal growth. These primary parameters will determine whether the novel RUTF ameliorates key symptoms of malnourishment in our MN model —reduced weight and length—and to what extent. After this first assessment, advanced analyses on micro- and molecular levels will evaluate the potential long-term effects of RUTF intervention. Bone analysis will assess microstructure and functionality, both of which are severely impaired by protein deficiency^19^. Optimised RUTF formulas are expected to improve factors like bone volume and mineralisation of the MN group, which should translate into enhanced mechanical strength and fracture resistance.

The MN group effectively models malnutrition on microbial and metabolomic levels, ensuring an accurate assessment of RUTF interventions. While alpha diversity did not significantly differ between CT and MN groups, distinct microbiome compositions were observed. For example, *Ruminococcus* species were less abundant in the MN group. These species are key indicators of gut health in children aged 6–24 months^9,41^ and are known to mitigate impaired growth phenotypes caused by undernourished microbiota^42^. Metabolomic analysis revealed distinctive profiles between CT and MN groups, with metabolites like acetate more abundant in the CT group. Acetate, produced by the gut microbiota, plays a protective role in maintaining the intestinal barrier, and altered acetate levels in the stool have been associated with diseases like ulcerative colitis^43^. Successful RUTF interventions should normalise microbiome composition and metabolomic profiles, which will be monitored through caecal analysis and stool sample time series.

This protocol provides a robust approach for RUTF intervention studies that go beyond traditional metrics of weight and length improvement. The MN group demonstrates a skeletal, microbial, and metabolomic profile representative of malnourishment, making it an ideal model for evaluating novel RUTF formulas. The holistic approach outlined here allows for predicting long-term outcomes beyond the study endpoint, offering critical insights to facilitate progression toward clinical trials.

The results generated through this protocol have significant implications for the design and production of improved RUTF formulations. The emphasis on local, cost-effective ingredients aligns with UNICEF’s guidelines^3^, promoting accessibility and sustainability. Moreover, insights into the interaction between RUTFs and the microbiome could guide the development of microbiota-directed complementary foods, addressing malnutrition and associated long-term health risks. These data can support regulatory approvals and the refinement of clinical trial designs, paving the way for broader adoption of optimised RUTFs in low-resource settings.

While this study provides a robust framework for evaluating RUTF efficacy, certain limitations must be acknowledged. First, using a rat model, while valuable for controlled mechanistic insights, may have limited translatability to human physiology and metabolism. Second, the controlled laboratory environment may only partially replicate the complex socio-environmental factors influencing malnutrition and recovery in diverse human populations. Third, the study primarily focuses on short-term recovery metrics, and while it incorporates markers predictive of long-term outcomes, extended validation in clinical contexts will be necessary. Additionally, the reliance on advanced multi-omics techniques, although offering comprehensive insights, presents challenges in scalability and cost for implementation in low-resource settings.

Nevertheless, these limitations are integral steps in the translational process, enabling the identification of key biomarkers and mechanisms that will guide subsequent human trials. By leveraging the insights gained from this preclinical model, future studies will address these gaps by testing novel RUTF formulations in diverse human populations under real-world conditions. This progression will ensure that the findings of this study are not only scientifically rigorous but also practically applicable, contributing to improved global malnutrition treatment strategies.

In conclusion, this study provides a rigorous framework for evaluating RUTF interventions, addressing critical limitations of current methods. Integrating skeletal, microbiome, and metabolomic analyses ensures a holistic approach to understanding malnutrition recovery, with significant potential to inform future clinical trials and global health policies. While challenges remain, the results from this study are expected to contribute substantially to advancing the treatment and prevention of severe acute malnutrition worldwide.

## Acknowledgements

We are grateful to Dr. Moses Mwangi for allowing us the use of the Equatorial Nuts Processors (ENP) laboratories to carry out the formulation of our novel RUTF. We thank Samuel Maina for providing the assistance of laboratory technicians to support the formulation process, Emma Mugo (team lead), Lorrine Akoth, Lucky Karubu, Charles Mburu, Brian Nalianya, Mary Murage, Florence Maina, and Lilian Gichuku with the participation of Benard Maguti (head of the laboratory), the Quality Assurance Manager. We are also grateful to Svetlana Penn, the laboratory technician and the students Mona Khalaf, Gal Becker, Aluma Haiman and Seman Daeem who helped support the animal study at HUJI. The opinions expressed in this paper do not necessarily reflect the views of our donors or partners.

